# Defining spatial relationships between spinal cord axons and blood vessels in hydrogel scaffolds

**DOI:** 10.1101/788349

**Authors:** Ahad M. Siddiqui, David Oswald, Sophia Papamichalopoulos, Domnhall Kelly, Priska Summer, Michael Polzin, Jeffrey Hakim, Bingkun Chen, Michael J. Yaszemski, Anthony J. Windebank, Nicolas N. Madigan

## Abstract

Positively charged oligo-polyethylene glycol fumarate (OPF+) hydrogel scaffolds, implanted into a complete transection spinal cord injury (SCI), facilitate a permissive regenerative environment and provide a platform for controlled observation of repair mechanisms. Axonal regeneration after SCI is critically dependent upon the availability of nutrients and oxygen from a newly formed blood supply. In this study, the objective was to investigate fundamental characteristics of revascularization in association with the ingrowth of axons into hydrogel scaffolds, and to define the spatial relationships between axons and the neovasculature. A novel combination of stereologic estimates and precision image analysis techniques are described to quantitate neurovascular regeneration in rats. Multichannel hydrogel scaffolds containing Matrigel-only (MG), Schwann cells (SCs), or SCs with rapamycin-eluting poly(lactic co-glycolic acid) (PLGA) microspheres (RAPA) were implanted for 6 weeks following complete spinal cord transection. Image analysis of 72 scaffold channels identified a total of 2,494 myelinated and 4,173 unmyelinated axons at 10 micron circumferential intervals centered around 708 individual blood vessel profiles. Blood vessel number, density, volume, diameter, inter-vessel distances, total vessel surface and cross-sectional areas, and radial diffusion distances in each group were measured. Axon number and density, blood vessel surface area, and vessel cross-sectional areas in the SC group exceeded that in the MG and RAPA groups. Axons were concentrated within a concentric radius of 200-250 microns from the blood vessel wall in Gaussian distributions which identified a peak axonal number (mean peak amplitude) corresponding to defined distances (mean peak distance) from each vessel. Axons were largely excluded from a 25 micron zone immediately adjacent to the vessel. Higher axonal densities correlated with smaller vessel cross-sectional areas. A statistical spatial algorithm was used to generate cumulative distribution F- and G-functions of axonal distribution in the reference channel space. Axons located around blood vessels were definitively organized as clusters and were not randomly distributed. By providing methods to quantify the axonal-vessel relationships, these results may refine spinal cord tissue engineering strategies to optimize the regeneration of complete neurovascular bundles in their relevant spatial relationships after SCI.

**Impact Statement:** Vascular disruption and impaired neovascularization contribute critically to the poor regenerative capacity of the spinal cord after injury. In this study, hydrogel scaffolds provide a detailed model system to investigate the regeneration of spinal cord axons as they directly associate with individual blood vessels, using novel methods to define their spatial relationships and the physiologic implications of that organization. These results refine future tissue-engineering strategies for spinal cord repair to optimize the re-development of complete neurovascular bundles in their relevant spatial architectures.

## Introduction

Traumatic spinal cord injury (SCI) results in motor, sensory and autonomic dysfunction due to neuronal and vascular disruption (1). Multichannel hydrogel scaffolds, implanted into a complete spinal cord transection model, allow for experimental manipulation of the local injury conditions. Hydrogels are a platform for the quantification (2–5) of repair mechanisms (6) in regenerating spinal cord tissue. Positively-charged hydrogel scaffolds fabricated from oligo(poly(ethylene glycol) fumarate) (OPF+) produce a permissive micro-environment for axonal regeneration (7, 8), and enhance neuron outgrowth density and directionality (9). Seeding OPF+ scaffolds with Schwann cells (SCs) provides neurotrophic and vasogenic signals as well as a peripheral nerve myelination structure around CNS axons (10). The multichannel scaffold adjoins specific anatomical tracts of the rat spinal cord, and enables independent measurements within scaffold channels as structurally separated areas (11, 12). OPF+ scaffolds containing SCs and embedded with rapamycin-releasing poly(lactic co-glycolic acid) (PLGA) microspheres reduced the foreign body response and promoted functional recovery after spinal cord transection (13). Rapamycin decreases inflammatory cell infiltration and activation in an SCI lesion environment (14, 15). The drug also reduces fibrosis in reaction to various implanted synthetic materials (16, 17). Rapamycin itself may enhance vascularization of regenerating tissue by normalizing blood vessel distribution (13, 22–25)

The availability of nutrients and oxygen from the regenerating blood supply is a critically important factor for axonal regrowth (18, 19). Vascular injury causes ischemia, hemorrhage, and increased permeability of vessels for the influx of inflammatory cells (20). Blood vessel beds which form after SCI are disorganized and function inefficiently (18). The relationships between regenerating axons and blood vessels as neurovascular bundles that form within hydrogel biomaterials are not well understood. A correlation between blood vessel diffusion distances, as a function of the distribution of small caliber blood vessels, and the number of regenerating axons, has been shown (21). The formation of a dense capillary structure with overlapping diffusion supply supported higher numbers of regenerating axons. A correlation between blood vessel formation and improvement of functional recovery after SCI has also been described (26–28).

In this present study, our objective was to quantify spatial relationships between the neovasculature and regenerating axons within hydrogel scaffolds using stereologic estimates and precision measurements by image quantification.

## Materials and Methods

OPF+ hydrogel scaffolds were implanted for six weeks into a complete transection injury with and without concomitant Schwann cell delivery and treatment with rapamycin. Materials and methods relating to PGLA microsphere and OPF+ hydrogel fabrication, SC culture, animal surgeries, scaffold implantation, tissue sectioning, antibodies, and immunohistochemistry have been published (12), and may be found in the Supplementary Data section.

### Image processing

Individual scaffold channels were imaged using a LSM510 laser scanning confocal microscope (Carl Zeiss, Inc., Oberkochen, Germany) at 20x magnification. 72 channels were imaged in OPF+ scaffolds with Matrigel and empty PLGA microspheres (n=5 animals, 16 channels); OPF+ scaffold with SCs and empty PLGA microspheres (n=6 animals, 34 channels); and OPF+ scaffolds with SCs and rapamycin-eluting PLGA microspheres (n=6 animals, 22 channels).

### Estimation of axons and blood vessels number, density and physiologic measures

Image analysis was performed by an investigator who was blinded to the animal group using Neurolucida software (version 11.062, MBF Bioscience, Williston, Vermont, USA). Core channel areas (21) were calculated from two orthogonal diameter measurements using the “Quick Measurement” Tool. Axons staining either with Tuj-1 (unmyelinated), or Tuj-1 co-localizing with MBP (myelinated) were digitally marked with separate symbols (Figure 1 A-B). The manual neuron-tracing tool was repurposed to outline blood vessels along their innermost surface after collagen IV staining. Numerical markers were assigned to each vessel (Figure 1 C-D). Blood vessel cross-sectional areas were calculated by the Neurolucida Explorer software. Total surface area of vessels was calculated as the summation of cross-sectional areas for all vessels within a given channel. A total of 708 blood vessel profiles, and 6667 axons (2,494 myelinated and 4,173 unmyelinated axons) were analyzed (Table 1).

**Figure 1:**
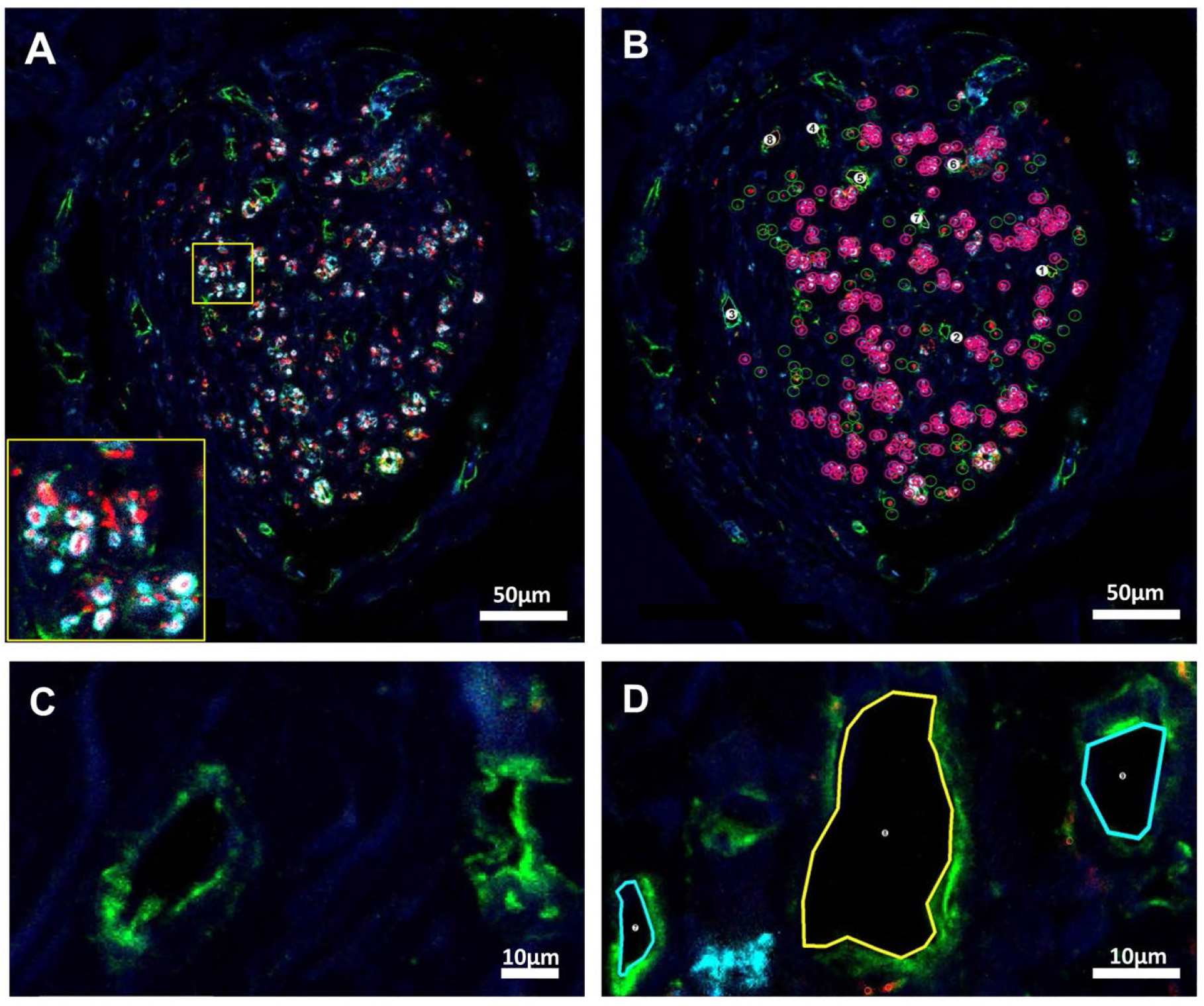
Image processing and marking strategies to identify and count axons and vessels and in OPF+ scaffold channels. **(A)** Confocal imaging of a Schwann cell (SC) channel within an OPF+ scaffold without rapamycin before marking. Scaffold channels were composed of an interior core of tissue contained regenerating axons and blood vessels, and a laminar, fibrotic outer layer adjacent to the scaffold channel which was relatively devoid of axons and vessel structures. Axons were labeled in transverse tissue sections with immunostaining for β-tubulin (red) and myelin basic protein (cyan). Unmyelinated axons appeared as punctate β-tubulin staining, while myelinated axons appeared as positive myelin basic protein immunostaining circumscribed around central axonal β-tubulin staining (insert). **(B)** Using Neurolucida software, manual marking designated unmyelinated axons with open green circles, myelinated axons with open pink circles and a central point, and blood vessels with numerical marks. Scale bar (A-B) = 50 μm. **(C)** Collagen IV staining outlined blood vessel walls (green). Vessels could be distinguished from cystic structures by the presence of a multi-layered wall containing identifiable endothelial cell nuclei and subendothelial collagen IV staining around an open lumen. **(D)** Vessels were outlined and numbered by the software. Scale bar (C-D) = 10 μm. DAPI staining (blue) identifies cell nuclei in all panels.

**Table 1:** Numbers of vessels and axons under analysis

| marked structures | vessels | um axons | m axons | total axons |
| --- | --- | --- | --- | --- |
| MG | 133 | 495 | 207 | 702 |
| SC | 346 | 2796 | 2218 | 5014 |
| RAPA | 229 | 882 | 69 | 951 |
| total | 708 | 4173 | 2494 | 6667 |

Estimates for blood vessel length density (Lv) were derived according to the formula Lv = 2QA where Q_A_ was the number of marked vessel profiles intersecting with tissue section plane within the measured area of the inner tissue channel core (30–32). Estimates for surface area density (Sv) of blood vessels were derived from direct measurement of each vessel luminal area. Estimates for blood vessel volume (Vv) were derived from luminal surface area measurements multiplied by the tissue section thickness (10 microns). Mean blood vessel diameter was calculated from the ratio of surface area to length densities, according to the equation:

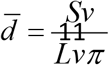

Radial diffusion distances were derived as an inverse proportion of the length density, according to the equation (30–32):

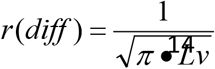

### Sholl analysis

Sholl analysis assessed the relationship between axon number and distance from blood vessels. A starting radius was defined for each vessel by manual measurement of its maximal diameter using “Quick Measurement” tool. The software then generated a series of concentric circles at interval distances of 10 μm. All myelinated and unmyelinated axons falling between the bounds of adjacent concentric circles were counted and their distance from the vessel center was recorded within a 10 μm bin. Manual measurement of the distances between all blood vessels was performed in replicate. The mean value of all inter-vessel distances was used as a cut-off to exclude axons which were expected to be too far away from the central blood vessel to be supplied by that vessel.

Axon counts were plotted for each radius interval using Prism’s Graph Pad software (La Jolla California USA). Each vessel was represented by an individual graph depicting unmyelinated, myelinated and total axon counts. A Gaussian function was fitted to each curve. 55 MG, 71 SC, and 65 RAPA channel vessels surrounded by respective totals of 59, 98, and 82 axons were excluded due to low axon numbers prohibiting an accurate Gaussian curve. High cumulative axon counts over the remaining vessels required a strategy to condense the datasets. Values for Mean Peak Amplitude for each blood vessel were derived as the y-axis number of axons at each Gaussian curve apex, and the Mean Peak Distance as the distance value on the x-axis corresponding to the amplitude apex. Peak densities of axons were calculated as the Mean Peak Amplitude divided by πr^2^, where r was defined as corresponding Mean Peak Distance.

### Cumulative Frequency Distribution Analysis

Immunofluorescence images of each channel were imported into Image J (National Institutes of Health, Bethesda, MD. (33)) using the Bio-Formats Importer to split the images into separate colors. The channel core area of reference was defined for the software by creating a manually outlined mask. A threshold was set to optimize contrast between the β-tubulin positive axons background. Thresholding parameters were held constant for each image undergoing analysis. The image was converted into a binary format. Cumulative Frequency Distributions of axon spatial relationships were generated using the Spatial Statistics 2D/3D Plug-in for Image J (34, 35). The plug-in generates curves of the percent frequency of points in an image (y-axis) that are located within defined distance of each other, from shortest to longest distance along the x-axis. The three statistical functions included the cumulative frequencies of the distances 1) between an axon and its nearest neighboring axon, termed the ‘G Function;’ 2) between an axon and a randomly generated point in the reference space, termed the ‘F Function;’ and 3) between 2 randomly generated points in the reference space, each with confidence interval calculated. Seven channels in each animal group were analyzed by the algorithm. The plugin generated x-y pairs at accumulating frequency intervals of 0.1% (1,000 data points) for the F or G function, and at 0.001 % intervals (100,000 data points) for the random curve generation. An Excel macro was therefore written to scan the dataset and pull each x-y value and confidence interval data at 1% cumulative frequency intervals, generating 100 data points per curve. Mean values of the x-values for each 1% frequency increment were calculated and plotted with confidence intervals.

### Statistics

Data are presented as mean values ± standard error of the mean, and confidence intervals are calculated for cumulative frequency distribution curves. One-way analysis of variance (ANOVA) with Tukey’s post hoc multiple comparisons test determined statistical significance between groups of means. Correlation analysis determined significance of paired relationships with two-tailed P values and Spearman r coefficients. p values of <0.05 were considered to be significant.

## Experiment

Following complete spinal cord transection, we investigated the spatial relationships of regenerating axons to the neovasculature through OPF+ scaffolds containing Matrigel only (MG) (n = 5 animals per group), Schwann cells only (SC) (n = 6), or SCs with sustained release of rapamycin (RAPA) (n = 6).

### Relationship of core area to axon numbers and density

Scaffold channels were seen to be composed of two compartments. An interior core of tissue contained regenerating axons and blood vessels. A densely laminar, fibrotic outer layer adjacent to the scaffold channel wall was devoid of axons and vessel structures (Figure 1A). Unmyelinated and myelinated axons were identified by immunohistochemical staining, and marked using the Neurolucida software (Figure 1A – B). Unmyelinated axons appeared as discrete points staining positively for β-tubulin (red), while myelinated axons appeared as points of β-tubulin circumscribed by myelin basic protein immunostaining (cyan). Blood vessels were identified by positive immunostaining for collagen IV (green) (Figure 1C). The internal lamina borders of each vessel were outlined (Figure 1D) to derive both vessel counts and lumen areas.

Myelinated and unmyelinated axons in MG, SC and RAPA channels were organized in small clusters (Figure 2 A-C). The mean total number of axons in SC channels (147.50 ± 17.03 axons/channel) was greater than that observed in MG (Figure 2D) (41.78 ± 8.35 axons/channel; p=<0.0001) or RAPA scaffold channels (43.23 ± 7.2 axons/channel; p=<0.0001; Figure 2D). SC channels supported higher numbers of both myelinated and unmyelinated axon subtypes. There were no differences in the mean number of blood vessels per channel between groups, which measured 8.31 ± 0.78 vessels in MG channels, 10.18 ± 0.62 vessels in SC channels and 10.41 ± 0.95 vessels in the RAPA group (Figure 2E).

**Figure 2:**
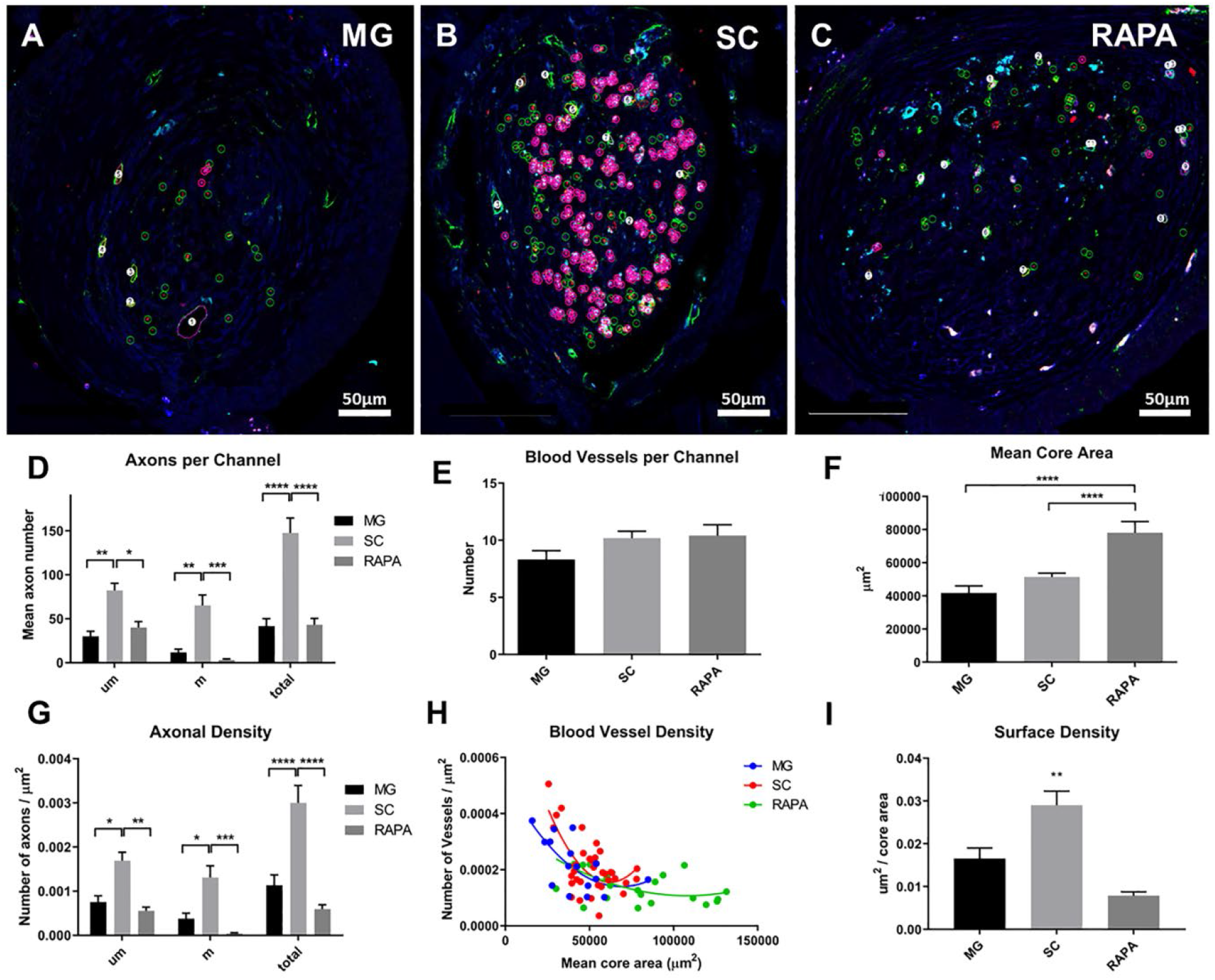
Axon and blood vessel number and density in relationship to core area. The core area, axon number, and blood vessel numbers were measured using Neurolucida software for rats implanted with OPF+ scaffold channels containing **(A)** Matrigel with sham PLGA microspheres (MG), **(B)** Schwann cells with sham microspheres (SC), or **(C)** Schwann cells with rapamycin microspheres (RAPA). Representative images of axonal densities with software markings are shown. Scale bar (A-C) = 50 μm. **(D)** Axon counts per scaffold channel (mean ± SEM) in each groups were calculated for unmyelinated (um), myelinated (m) and total axons. **(E)** Vessel number (mean ± SEM) per channel in each animal group. **(F)** Interior core areas mean ± SEM of MG, SC and RAPA channels. **(G)** Axonal densities were defined as the number of axons per core area (μm2), were calculated as mean ± SEM for unmyelinated (um), myelinated (m) and total axons. **(H)** Blood vessel densities demonstrated a negative correlation between mean core area and vessel density for all groups (MG: Spearman r=-0.5795, p=0.0186; SC: Spearman r=- 0.5129, p=0.0019; RAPA: Spearman r=-0.4884, p=0.0211). **(I)** Surface densities (Sv) of blood vessels as mean ± SEM per channel. * p<0.05, **p<0.01, ***p<0.001, ****p<0.0001.

**Figure 3:**
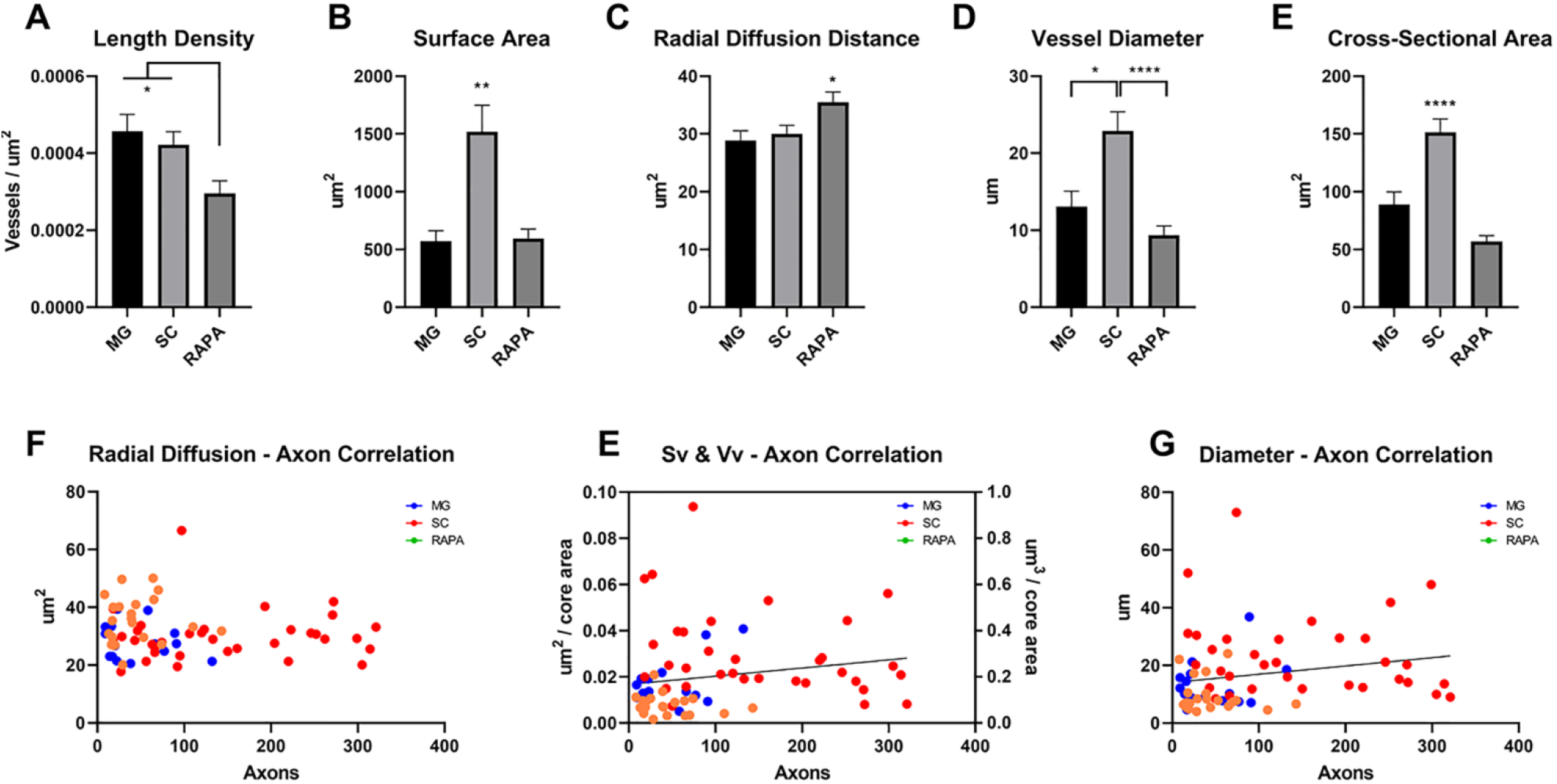
Blood vessel physiologic parameters in relation to axon number. **(A)** Length density (Lv) of blood vessels as a value that represents the combined length of all vessel profiles within a channel section plane was calculated as mean ± SEM for each group. **(B)** The surface area coverage of blood vessels per channel was the summation of each individual cross-sectional area measurement (mean ± SEM). **(C)** The radial diffusion distance was calculated as the inverse proportion of the Lv measurements per channel. Radial diffusion distance defines a cylindrical zone of nutrient and blood gas diffusion around the vessel wall (30, 31). **(D)** Blood vessel diameters were derived from stereologic estimates of the ratio of Lv and Sv. **(E)** Mean ± SEM cross-sectional areas of blood vessels in each channel type, by direct measurement of lumen area. **(F)** Mean radial diffusion distances did not correlate with total channel axonal number for a given channel, but ranged consistently between 20 and 40 μm^2^ with increasing axon numbers supported. **(F)** Positive correlations between axonal number were shown between Sv and the volume densities (Vv, which equals Sv multiplied by the thickness of the tissue section, 10 μm) (Spearman coefficient = 0.3217, p=0.006). **(G)** Positive correlations were also observed between axon number and increasing blood vessel diameters (Spearman coefficient = 0.2716, p=0.022). * p<0.05, **p<0.01, ****p<0.0001.

Axons and blood vessels in the RAPA group were distributed across a larger mean core channel inner surface area (78,042 ± 6,817 μm^2^) than the MG (41,829 ± 4,175 μm^2^) (p=<0.0001) and SC (51,407 ± 2,272 μm^2^) (p=0.0013) group (Figure 2F). This observation was consistent with an anti-fibrotic effect of rapamycin in scaffold channels in allowing for an expansion the inner tissue lumen (13). The mean density of total, myelinated, and unmyelinated axons, as the number of axons per μm^2^ of core surface area, was more concentrated (0.0030 ± 0.0004 total axons/μm^2^) in SC-seeded channels than in MG (0.0012 ± 0.0002 total axons/μm^2^; p=<0.0001) or RAPA groups (0.0006 ± 0.0001 axons/μm^2^; p=<0.0001; Figure 2G). Greater core tissue areas in the RAPA group produced longer distances between individual blood vessel profiles. The mean inter-vessel distance in the RAPA group (146.1 ± 8.65 μm) was higher than vessel distances in the MG group (96.94 ± 5.66 μm; p=<0.0001) and SC (116.5 ± 3.947 μm; p=0.0013. There was a negative correlation between vessel density and core area in all three groups (MG: Spearman r=- 0.5795, p=0.0186; SC: Spearman r=-0.5129, p=0.0019; RAPA: Spearman r=-0.4884, p=0.0211; Figure 2H). Blood vessel densities as a function of mean core area were lowest in the RAPA group, as shown by the rightward shift of the RAPA correlation curve. The surface area densities (Sv) of blood vessels in SC channels averaged 0.029 ± 0.003 μm^2^ per core area, greater than in MG (0.017 ± 0.002 μm^2^ per core area) (p<0.01) and RAPA channels (0.008 ± 0.001 μm^2^ per core area) (p<0.01) (Figure 2I).

### Analysis of blood vessel formation in OPF+ scaffold channels

Analysis of blood vessels quantitated vessel distribution, morphology and radial diffusion distance. The length density of vessels is a value that represents the combined length of all vessel profiles within a channel section plane. The mean length density (Lv) of blood vessels was calculated to be lower in RAPA channels (3.0 x 10^−4^ vessels/μm^2^) than MG (4.5 x 10^−4^ vessels/μm^2^) (p<0.05) and SC (4.2 x 10^−4^ vessels/μm^2^) (p<0.05) (Figure 3A). The blood vessel surface area was larger in the SC group, with vessels occupying a mean area of 1519.0 ± 229.0 μm^2^ per channel, than in MG (572.0 ± 89.8 μm^2^ or RAPA channels (594.3 ± 83.90 μm^2^) (Figure 3B). Blood vessel volume measurement followed the same trends, with far larger mean vessel volumes of 1.519 x 10^5^ ± 0.230 x 10^5^ μm^3^ per channel in the SC group as compared with 5.720 x 10^4^ ± 8.98 x 10^4^ μm^3^ in MG (p<0.01) and 5.943 x 10^4^ ± 0.84 x 10^4^ in RAPA channels (p<0.01).

Additional physiologic parameters were derived from measurements of the Lv and Sv. The radial diffusion coefficient, a measure of a cylindrical zone of diffusion around a blood vessel, was calculated as the inverse function of the length density (30, 31) The diffusion distance for Schwann cell vessels was 30.00 ± 1.48 μm^2^, 28.89 ± 1.64 μm^2^ in MG vessels and 35.52 ± 1.72 μm^2^ (p<0.05) in RAPA channels (Figure 3C). Mean blood vessel diameters per channel were calculated from the ratio of the Lv to Sv. The mean diameter of blood vessels in SC channels was 22.90 ± 2.46 μm, was greater than the mean vessel diameters in MG vessels (13.10 ± 2.00) (p<0.05) and in RAPA vessels (9.36 ± 1.21 μm) (p<0.0001) (Figure 3D). This stereologic calculation of diameter was consistent with direct measurements of cross-sectional areas of individual blood vessels in channels. The mean cross-sectional area per blood vessel in SC channels was 151.40 ±11.45, larger than that measured in MG (88.90 ± 11.08 μm^2^) and RAPA vessels (74.57 ± 4.93 μm^2^)(both p<0.0001) (Figure 3E).

Axon counts trended to be lower in the RAPA group with higher diffusion distances (Figure 3E) but correlations were not statistically significant. Diffusion distances consistently ranged between 20 μm and 50 μm with increasing axon number. The number of axons regenerating had positive correlations to the surface and volume area densities of vessels (Spearman coefficient = 0.3217, p=0.006) (Figure 3F) and to the diameter of blood vessels (Spearman coefficient = 0.2716, p=0.022) (Figure 3G).

### Spatial relationships of regenerating axons to blood vessels in scaffold channels

To quantitate the spatial distribution of single myelinated and unmyelinated axons around individual blood vessels, a novel methodology was developed around a Sholl sub-analysis. Concentric rings were centered upon each blood vessel increased in diameter by 10 μm intervals. Myelinated and unmyelinated were counted within the boundaries of each concentric ring interval (Figure 4A, centering on Vessel 9, for example). The axonal number in each ring interval was plotted as a distribution function against their distances from the blood vessel.

**Figure 4:**
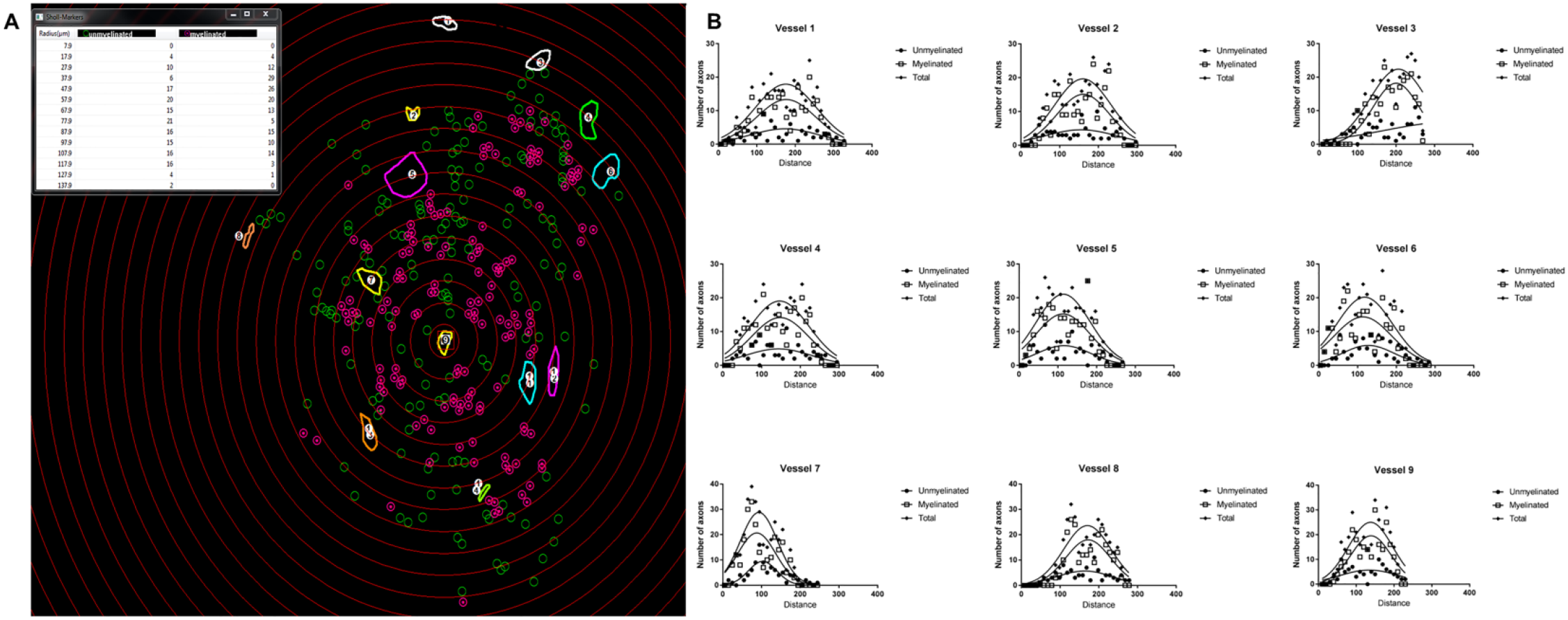
Sholl analysis Gaussian distributions of axons number with distance around blood vessels. Measurement of axon distribution was done using a Sholl analysis. Concentric rings were generated by the software in 10 μm intervals around each vessel, here Vessel 9. The rings were centered following determination of the vessel diameter as the innermost ring. Measured diameters ranged from 3.3 to 67.3 μm. Counts for unmyelinated (open circles) and myelinated axons (encircled points) which were located between two adjacent rings were tabulated by the software. **(B)** The number of unmyelinated, myelinated and total axons within each 10 micron radial increment was plotted as a function of the distance from each blood vessel, and consistently demonstrated a Gaussian distribution. This data was compressed by using the value of Mean Peak Amplitude to sample axon number at the maximal height of the distribution curve, and Mean Peak Distance as the corresponding distance from the vessel at which the distribution was maximal.

Axonal distributions around blood vessels were observed to be Gaussian (Figure 4B). Since the number of individual axons counted around each vessel within a given channel was cumulatively high, the dataset was condensed into values for Mean Peak Amplitude, as the number of axons on the y-axis represented by the peak of each Gaussian curve. The Mean Peak Distance was the distance on the x-axis at which the axonal number / amplitude was at its peak.

Plotting mean peak amplitude against mean peak distance (Figure 5) identified that the distance of maximal axonal number from a vessel was located within a radius of less than 200 μm from the vessel wall in the MG (Figure 5A). The radius for maximal axonal number extended to 250 μm in the SC groups and RAPA group (Figure 5B-C). The maximal number of axons was essentially excluded from a 25-30 μm radius immediately adjacent to a blood vessel, as observed in each animal group and whether the axons were myelinated, or unmyelinated (Figure 5). This effect was slightly less pronounced in the RAPA group, where two peak amplitudes of representing less than 4 unmyelinated axons were located within the exclusion zone.

**Figure 5:**
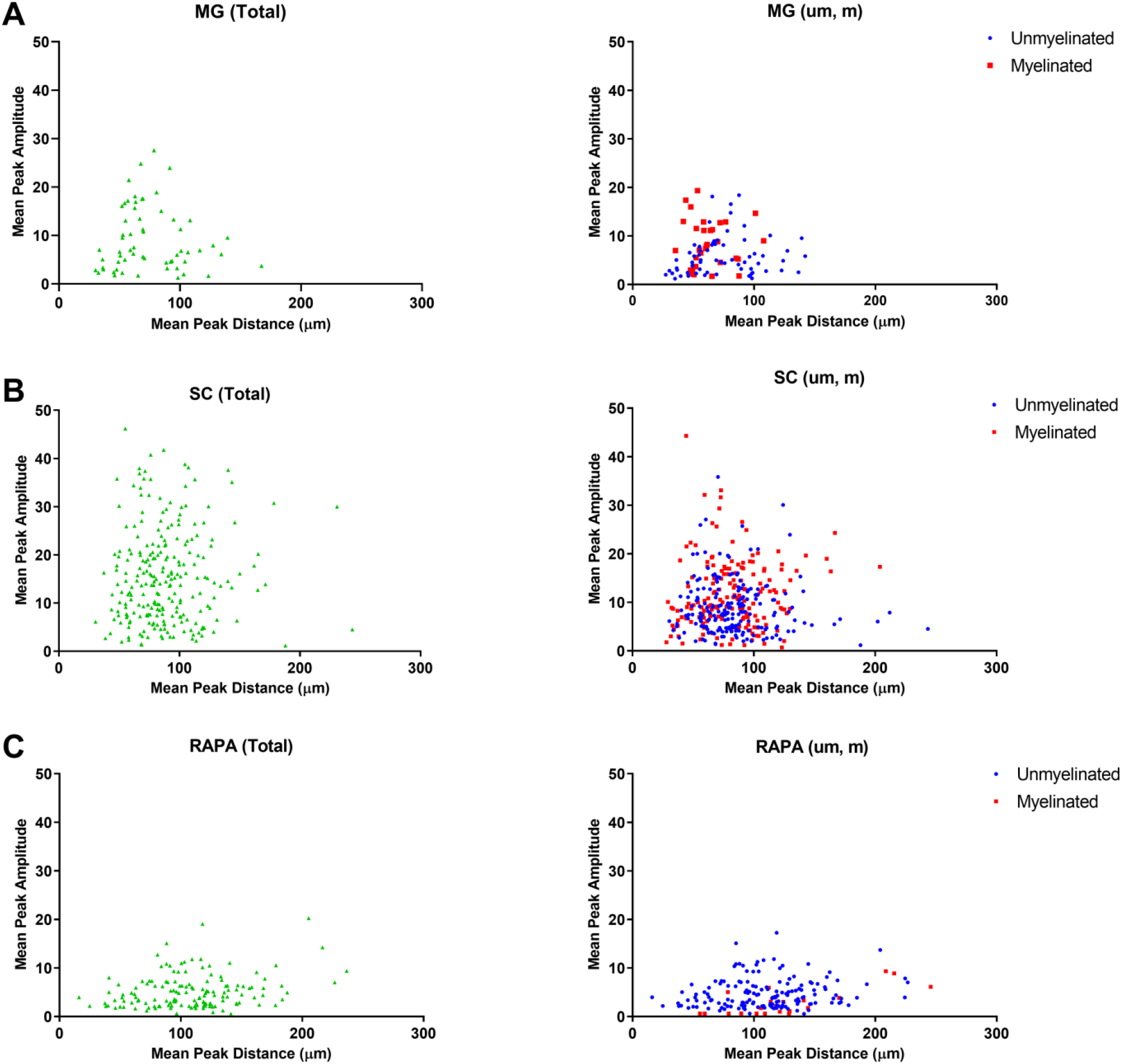
Mean peak amplitude and mean peak distance of axons compared to blood vessel cross sectional area. Total, unmyelinated (um), and myelinated (m) counts at Mean Peak Amplitude were plotted against their respective Mean Peak Distances in (A) MG channels, (B) SC channels, and (G-I) RAPA channels. These relationships identified a zone of exclusion of 25-30 μm immediately adjacent to the blood vessel wall where axons were not located at their Mean Peak Amplitudes.

For each blood vessel, the mean peak amplitudes for axonal concentrations were described as a function of the cross-sectional areas of the blood vessel around which the axons were distributed. Peak axonal density represented the peak amplitude of axons with respect to the circular area of intervening tissue determined by the mean peak distance as the radius from the vessel wall. For SC channels (Figure 6A), significant negative correlations were shown between peak axonal density of total axon amplitudes and vessel cross-sectional area for total (Spearman r=-0.1864, p=0.0035), unmyelinated (Spearman r=−0.1473, p=0.0252) and myelinated axons (Spearman r=- 0.2023, p=0.0074) axon distributions (Figure 6A). Correlations were not identified in MG or RAPA channels where the axonal densities lower (Figure 6 B and C).

**Figure 6:**
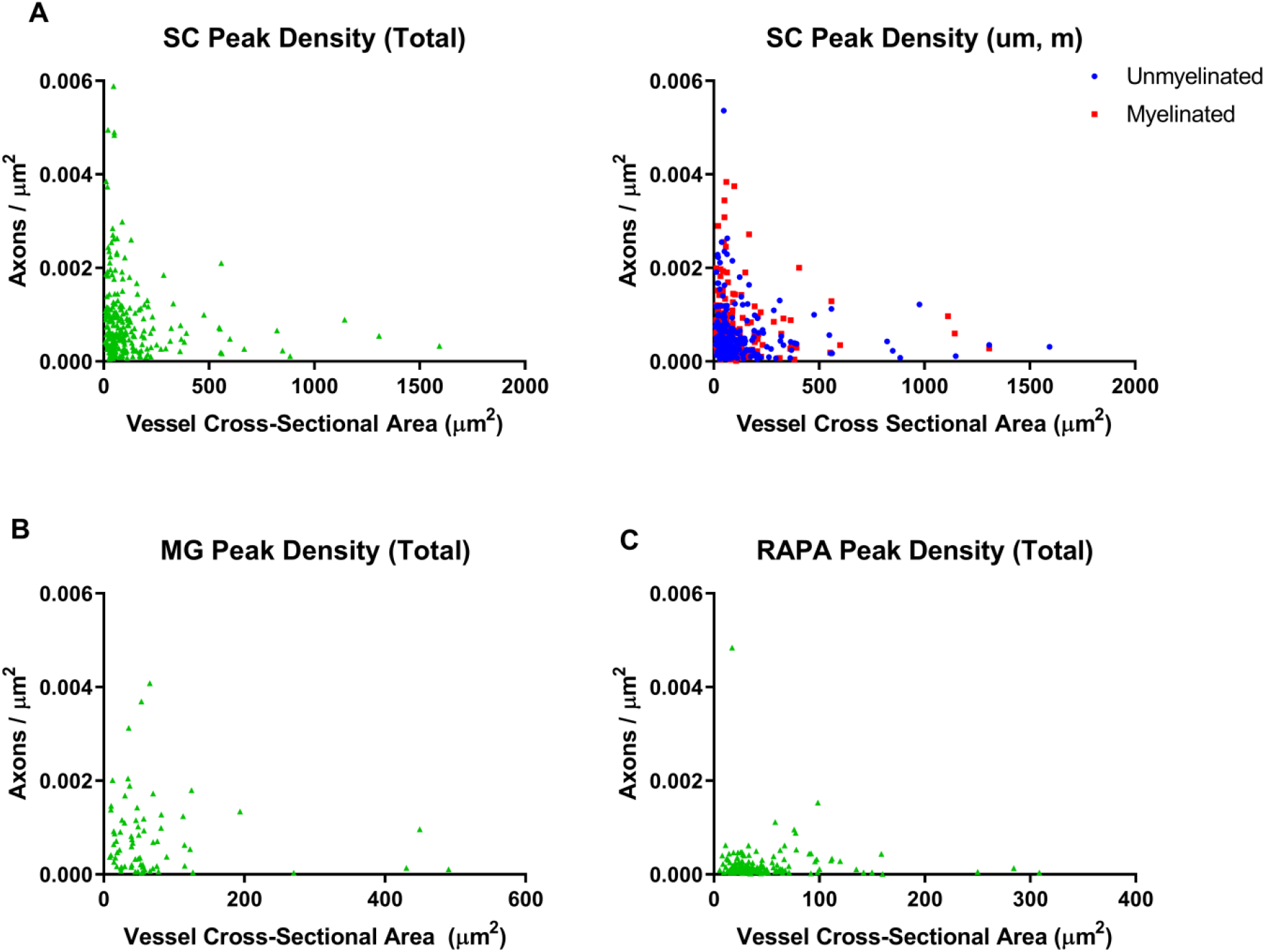
Relationship of axonal density to vessel cross-sectional area. Peak axonal density was derived as the Mean Peak Amplitude within the surface area defined by the Mean Peak Distance as the radius. Peak axonal density for each vessel were plotted to correlate with the vessel cross-sectional area of the reference vessel in **(A)** SC, **(B)** MG and **(C)** RAPA group channels. A negative correlation was observed in the SC channels, total (Spearman r=-0.1864, p=0.0035), unmyelinated (Spearman r=-0.1473, p=0.0252) and myelinated axons (Spearman r=-0.2023, p=0.0074) axon distributions. Density - vessel area correlations were not identified in MG or RAPA channels.

### Distribution and relationship of regenerating axons to each other

Cumulative distribution functions of inter-axon distances were generated using the Spatial Statistics 2D/3D Plug-in for NIH Image J (34, 35) (Figure 7). The distribution of axons within regenerating tissue could be considered to be random, clustered, or dispersed points in space (Figure 7A-C). Cumulative frequencies of the percentage of axons located within a given distance from another axon defined the G-function, or a nearest neighbor analysis. The G-function assessed whether axons were distributed in clusters or were dispersed relative to the nearest neighbor analysis of randomly generated points. In each animal group, the shift in the cumulative distribution G-curves of axon distances far to the left of the random point distribution curve signified a grouped relationship between axons (Figure 7D-F). The closest neighbor analysis also demonstrated that 90% of axons in the MG or SC groups were each located under 10.1 μm (CI 9.1 − 11.2 μm) and CI 8.8 − 11.3 μm) respectively from the next closest axon (Figure 7D and 7E). The RAPA group (Figure 6F) was slightly more dispersed with 90% of the axons being located within 20.6 μm (CI 24.2 – 32.4 μm) from the next closest axon.

**Figure 7:**
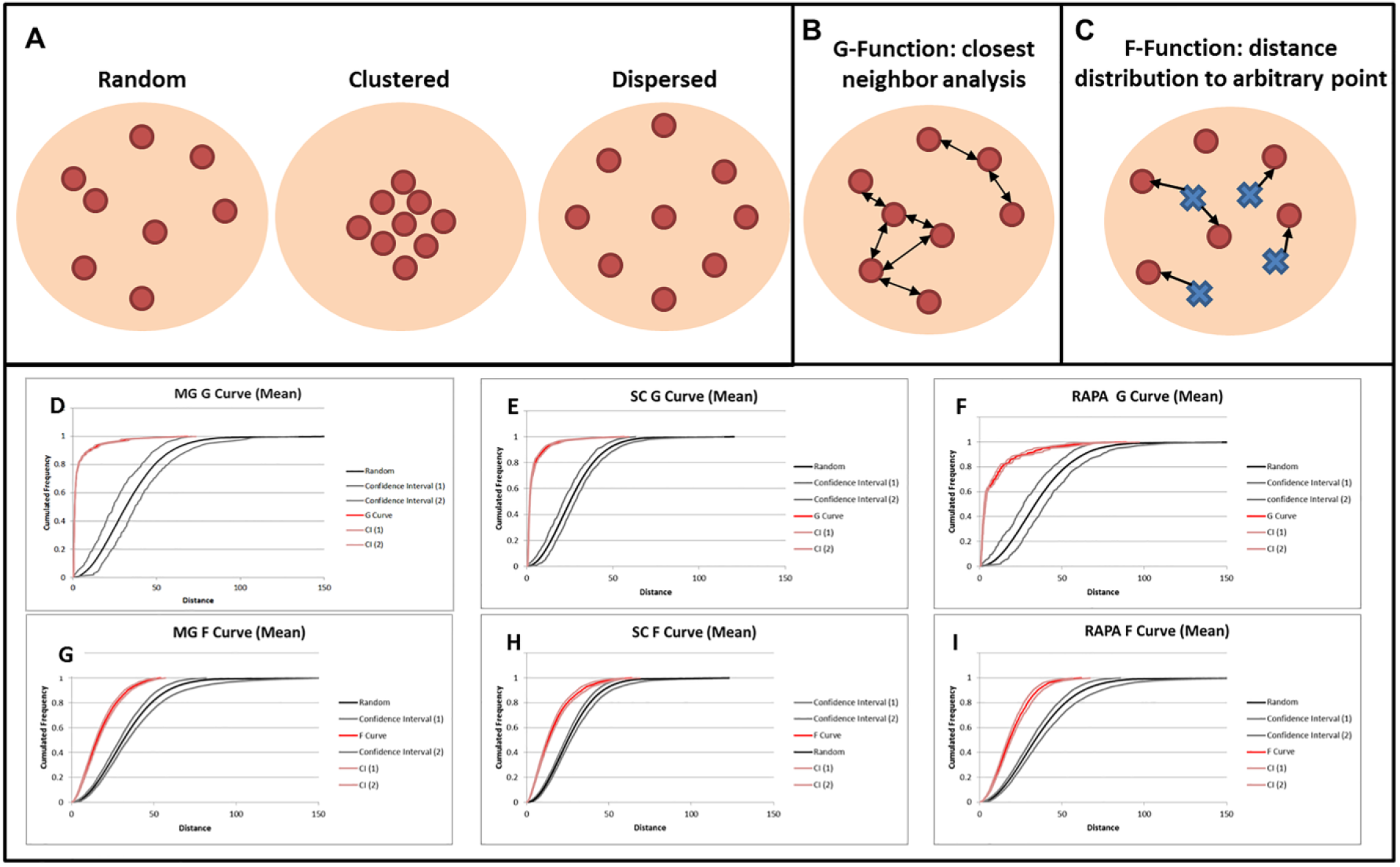
Analysis of axon clustering. **(A)** Axon distributions can be classified as random, clustered, or dispersed. **(B)** The closest neighbor analysis (G-function) measures the cumulative distance distributions between axons. **(C)** The F-function measures the distance distributions between an axon and a randomly generated point within the reference space. **(A-C)** are adapted from reference (35). The cumulative frequency of the percentage of axons or points (y-axis) at increasing distances from each other (x-axis) are depicted as G-curves, F-curves and random point curves confidence intervals for MG **(D, G)**, SC **(E, H)** and RAPA **(F, I)** channels. A grouped or clustered distribution of axons is demonstrated by the leftward shift of the F- or G- curve from the reference curves of random point distances.

Cumulative frequencies of the distances between an axon and a randomly generated point in the reference space were defined as the F-function. This analysis determined whether axons were distributed randomly, in definitive clusters or in a dispersed organization. The shift in each F-function curve to the left of a curve of distances between random points demonstrated that the relationship between regenerating axons in each animal group was in a definitely clustered, and non-random distribution (Figure 7 G-I). The F-curve shift signified that the distance between an axon and a random point was consistently shorter than the distance between two randomly generated points, indicating that axons lie closer together than if they were mathematically distributed in a random pattern. For example, 90 % of MG channel axons would be located within 34.0 μm of a random point (CI 30.6 – 34.5 μm), as compared to 90% of random points being at within a distance of 58.5 μm (CI 52.4 – 68.3 μm).

## Discussion

In this study, novel methods are described by which axonal regeneration and revascularization can be quantified in their innate spatial relationships. This approach adds to and can validate methods of stereologic estimation (30, 32) by means of systematic and detailed measurements of the distribution of single axons in relation individual blood vessels. In contrast to differential support of axonal regeneration between groups, blood vessel neovascularization occurred with similar numbers of vessels over time in each condition. Blood vessels in channel areas which were less fibrotic following rapamycin treatment needed to influence larger areas of core tissue, with implications for vessel morphology, tissue oxygenation and nutrient distribution. SC channel blood vessel coverage was more extensive, and the vessel caliber larger, than MG and RAPA groups. Smaller vessels within that neovasculature supported improved regeneration of axonal densities. By multiple measures, including surface area coverage, vessel volume and blood vessel diameter, significantly more blood could be delivered to SC channels in association with improved radial diffusion distances and higher axonal counts. Axonal growth was seen to observe a given distance from the vessel wall as opposed to regenerating in very close approximation to the vessel wall. This finding may relate to radial diffusion physiology with peak axon numbers at optimal positions in nutrient and gas diffusion gradients. Plotting the peak axonal density values against vessel cross-sectional area confirmed a correlation between vascular caliber and support of axonal regeneration. This results supports our previous observations that that improved axonal regeneration correlated with smaller radial diffusion distances and redundancy of vessel diffusion overlap (21).

Administration of rapamycin resulted in improved functional recovery following spinal cord transection in our previous study (13). Rapamycin is an allosteric mTOR inhibitor which has been shown to aid axonal regeneration following CNS and PNS injuries (36–39). Treatment with rapamycin and SCs here produced fewer regenerating axons than SCs alone. Our study importantly identified this phenomenon, but its scope was limited in that an explanation for this was not explored. Improved motor function recovery does not solely depend on axon density, but also may be associated with reduced fibrosis and inflammation (40–42). Rapamycin treatment has decreased cytokine production, and activation of microglia with functional improvements after SCI (13, 41). Rapamycin may have directly reduced the number of supporting SCs in our scaffold channels due to an anti-proliferative effect. Rapamycin inhibits the proliferation of fibroblasts and increases apoptosis (43). Reduction in the number of SCs relative may in turn impair their angiogenic and neurotrophic support. Additional studies using rapamycin in SCI, have identified an anti-angiogenic effect (44–46).

Micro-vessels in the CNS provide trophic support (47–49), and regenerating axons have been shown to grow along blood vessels (50). Strategies to limit vascular damage and restore blood flow to the injured cord could enhance spinal cord repair (51, 52). Increased blood vessel density correlates with improvements in neurologic function after spinal cord injury (26–28). The anatomic distribution of axons and blood vessels is critical in neurodevelopment, and ‘re-development’ of the spinal cord after injury is an important tissue-engineering goal. Neuropilin 1 (NRP1) knock-out in endothelial cells in a mouse model produced abnormal formation of both blood vessels and axonal distribution (53). Vessel diameter was larger in the *Nrp1^fl/−^; Tie2-Cre* mutants, suggesting that smaller blood vessels facilitate normal axon formation.

Axons around blood vessels regenerated in defined clusters in OPF+ channels, which may represent axonal sprouting or multiple individual axons in parallel groups. In development, axons distributing from lateral geniculate nucleus follow chemical and electrical cues (54), in that axons that are the nearest to each other are electrically active at the same time. This synchrony of activity may help axons navigate to a target in groups. Blocking this electrical activity with tetrodotoxin led to abnormal patterning of connections to the visual cortex (55). The closer proximity and greater number of axons could aid in pathfinding of spinal axons in OPF+ scaffolds.

## Conclusion

Hydrogel scaffolds have provided a detailed model system to investigate the regeneration of axons and blood vessels after SCI, using novel methods to define their spatial relationships. Key observations include that SCs within hydrogel channels supported superior neurovascular bundle regeneration than in the MG or RAPA groups, in axon and vessel density, and in physiologic parameters of vessel diameter, and radial diffusion distances. Neurovascular bundles are represented spatially by a Gaussian distributions of axons around centralized vessels, with an area of relative exclusion of axonal regeneration in a 25-30 μm immediately adjacent to the vessel wall. Important correlations for improving axonal growth include surface area densities of vessels and smaller vessel cross-sectional areas. Axons regenerate in spatial clusters defined mathematically by cumulative distribution functions. These results may refine future tissue-engineering strategies for SCI repair to optimize the regeneration of complete neurovascular bundles in their relevant spatial architectures.

## Acknowledgments

The authors would like to acknowledge the technical assistance of Jarred Nesbitt and administrative assistance of Jane Meyer.

## Author Disclosures and Conflicts of Interest

### Author disclosure statements

Dr. Siddiqui reports no disclosures or conflicts of interest.

Dr. Oswald reports no disclosures or conflicts of interest.

Dr. Papamichalopoulos reports no disclosures or conflicts of interest.

Mr. Kelly reports no disclosures or conflicts of interest.

Dr. Summer reports no disclosures or conflicts of interest.

Mr. Polzin reports no disclosures or conflicts of interest.

Dr. Hakim reports no disclosures or conflicts of interest.

Dr. Chen reports no disclosures or conflicts of interest.

Dr. Yaszemski reports no disclosures or conflicts of interest.

Dr. Windebank reports no disclosures or conflicts of interest.

Dr. Madigan reports no disclosures or conflicts of interest.

### Funding

This work was generously funded by grants from the National Institute of Biomedical Imagining and Bioengineering (EB02390), National Institute of Biomedical Imagining and Bioengineering (TL1 TR002380), Morton Cure Paralysis Fund, and Craig H. Nielsen Foundation.

## Supplementary Data

### Animals

The study was approved by the Mayo Clinic Institutional Animal Care and Use Committee (IACUC). 17 adult female Fischer rats (approximately 200 g; Harlan Laboratories, Indianapolis, USA) were held in conventional housing on a 12 hour light-dark cycle with free access to water and chow. Care by technicians and veterinarians experienced in management of rat spinal cord injury were available daily without interruption.

### Poly-lactic-co-glycolic acid (PLGA) microsphere fabrication

Drug-eluting microspheres were fabricated using a water-in-oil-in-water double emulsion and solvent evaporation technique, as we have previously described (13). Briefly, 250 mg of 50:50 D,L-poly-lactic-co-glycolic acid (PLGA) (50:50 DLG 4A, molecular weight 29 kDa, Evonik Industries AG, Essen, Germany) were dissolved in 1 ml of methylene chloride. 100 μl of a 10 mg/ml solution of rapamycin (Toronto Research Chemicals, Toronto, Canada) in absolute ethanol, or ethanol vehicle (empty microspheres) were added and emulsified by vortexing. Two ml of 2% poly(vinyl alcohol) (PVA) (Sigma-Aldrich, St. Louis, MO, USA) were added dropwise while vortexing for 30 seconds, and the resulting microsphere suspension was then poured into 100 ml of 0.3% PVA under gentle stirring on a magnetic stir plate. After one minute, 100 ml of 2% isopropyl alcohol was added and the suspension was left under gentle stirring for one hour to evaporate off the methylene chloride. The PLGA microsphere suspension was then centrifuged at 2500 RPM and washed four times with distilled water. The microspheres were placed in a −80° C freezer overnight, and freeze-dried under high vacuum.

### OPF+ scaffold fabrication

Synthesis of OPF macromere was performed via condensation reaction between polyethylene glycol (PEG) and triethyamine, as previously described (29). 1g of OPF powder was dissolved in 650μl of deionized water, 0.05% (w/w) of photoinitiator (Irgacure 2959, Ciba Specialty Chemicals, Tarrytown, New York, USA) and 0.3 g of N-vinyl pyrrolidinone, a cross-linking reagent. Chemical modification with the positively charged monomer [2- (methacryloyloxy)ethyl]-trimethylammonium chloride (MAETAC) (80% wt in water; Sigma-Aldrich) followed at 20% w/w (9), producing OPF+ polymer solution. 25 mg of PLGA microspheres were added to 250 ul OPF+ liquid polymer thereafter. Scaffold fabrication was as described previously (7) via injection of the liquid polymer with suspended microspheres into a tubular glass mold containing seven parallel aligned wires as placeholders for the channel spaces. The casting was polymerized by 365 nm UV light exposure for one hour. Upon rehydration in PBS, cylindrical segments were cut to lengths of 2 mm, yielding scaffolds that were 2.6 mm in diameter and had seven longitudinal channels of 450μm diameter. The release kinetics of rapamycin microspheres embedded in OPF+ scaffolds, and the *in vitro* and *in vivo* physiologic effect of drug release, have been characterized in our model (13).

### Schwann Cell (SC) isolation and culture

SCs were cultured from the sciatic nerve of 2-5 day old rat pups, as we have previously described (7). Cells were grown in SC media (DMEM/F12 medium containing 10% fetal bovine serum, 100 units/mL penicillin/streptomycin (Gibco, Grand Island, New York, USA), 2 μM forskolin (Sigma-Aldrich), and 10 ng/mL recombinant human neuregulin-1-β1 extracellular domain (R&D Systems, Minneapolis, Minnesota, USA)). SCs were initially plated onto 35 mm laminin-coated dishes and incubated at 37° C in 5% CO2 for 48 hours, and then transferred to tissue culture flasks for expansion over 4 passages before use in scaffolds.

### OPF+ scaffold loading and surgical scaffold implantation

Scaffolds were sterilized by immersion in serial dilutions of ethanol (80%, 40%, 20% for 20 minutes each), followed by two washes in distilled water and incubation in SC media. OPF+ scaffolds with empty PLGA microspheres were loaded with SCs or Matrigel (MG) alone. OPF+ scaffolds with rapamycin releasing microspheres were loaded with SCs (RAPA), as previously described (6, 10, 13). SC and RAPA scaffolds contained SCs resuspended in 8 ul of pre-chilled Matrigel (BD Biosciences, San Jose, California, USA) at a density of 10^5^ cells/μl. Loading was performed using a gel-loading pipette tip under microscopic view at 4°C temperature, followed by three minutes at 37°C. Scaffolds were incubated in SC media overnight prior to use in animal surgeries. Animal spinal cord transection surgical techniques, scaffold implantation and post-operative care were as we have previously described (8). Anesthesia was performed via intraperitoneal injection of a ketamine (80 mg/kg; Fort Dodge Animal Health, Fort Dodge, Iowa, USA) and xylazine (5 mg/kg; Lloyd Laboratories, Shenandoah, Iowa, USA). Laminectomy through the T8-T10 level was followed by complete T9 spinal cord transection. OPF+ scaffolds were implanted within a 2 mm gap between the retracted spinal cord stumps, according to the experimental groups, and the surgical wound was closed in anatomical layers.

### Tissue preparation and sectioning

Six weeks post-surgery, animals were sacrificed by intraperitoneal injection of 0.4 mL sodium pentobarbital (40 mg/kg) (Fort Dodge Animal Health, Fort Dodge, Iowa, USA) and transcardial perfusion with 4% paraformaldehyde in PBS. The vertebral column with spinal cord was removed *en block* and post fixed overnight in 4% paraformaldehyde at 4° C. The lesion site with scaffold and adjacent spinal cord ends were carefully dissected free from the vertebral column in 15 mm segments and processed for paraffin embedding. Segments of spinal cord containing the scaffold were subsequently cut into 10 μm transverse sections on a Reichert-Jung Biocut microtome (Leica, Bannockburn, Illinois, USA) and collected on numbered slides. Tissue sections from the midpoint of the scaffold length were selected for subsequent immunohistochemistry analysis of individual channels.

### Antibodies and immunohistochemistry

Primary antibodies included Tuj-1 (βIII-tubulin) for the identification of axons (mouse anti-rat, 1:300, Millipore Chemicon, Temecula CA USA); myelin basic protein (MBP) for myelinated axons (goat anti-rat, 1:400, Santa Cruz Biotechnologies, Dallas, TX USA); and collagen IV for the blood vessel basement membrane (rabbit anti-rat, 1:800, Abcam, Cambridge, Massachussets, USA). Secondary antibodies included Cy3-conjugated affinity purified donkey anti-mouse IgG (1:200, Millipore Chemicon) (B-III tubulin); AlexaFluor™ 647-conjugated AffiniPure donkey-anti goat IgG (1:200, Jackson ImmunoResearch Laboratories, West Grove, Pennsylvania, USA) (MBP); and Cy2-conjugated donkey anti-rabbit IgG antibody (1:200, Millipore).

Slides were deparaffinized in xylene and rehydrated by serial immersion in graded ethanol (100%, 95%, 70%) and distilled water for 5 minutes each. Antigen retrieval was performed by immersion in 1 mM EDTA in PBS, pH 8.0 in a steamer for 30 minutes (97 degrees C). Sections were washed with PBS with 0.1% Triton X-100, and blocked with 10% normal donkey serum in PBS with 0.1% Triton X-100 for 30 minutes. Primary and secondary antibodies were diluted in PBS with 5% normal donkey serum and 0.3% Triton X-100. Sections were incubated with primary antibodies at 4 degrees C overnight and secondary antibodies for 1 hour following washing. Tissue was mounted under glass coverslips using Slow Fade Gold Antifade Reagent with DAPI nuclear stain (Molecular Probes, Eugene, Oregon, USA).

## Notes

### Competing Interest Statement

The authors have declared no competing interest.

### Summary of Updates

This version includes new data, updated figures, and additional interpretations.

